# A-Islands: A plant dataset for biodiversity research and species monitoring on Australian islands

**DOI:** 10.1101/2024.08.29.609992

**Authors:** Julian Schrader, David Coleman, Ian Abbott, Sally Bryant, Ralf Buckley, Darren Crayn, Rachael V. Gallagher, Stephen Harris, Harold Heatwole, Betsy Jackes, Holger Kreft, Kevin Mills, Jamie Kirkpatrick, Peter K. Latz, John Neldner, Cornelia Sattler, Micah Visoiu, Elizabeth H. Wenk, John C. Z. Woinarski, Stuart Worboys, Ian J. Wright, Isabel Zorn, Mark Westoby

## Abstract

Australia’s coastline is fringed by more than 8,000 continental islands. These islands feature a diverse array of landforms, rock and soil types and geological origins. Some of these islands are among the least invaded, most pristine habitats in Australia and support high plant diversity. Here, we present a new Australia-wide curated dataset for plant species occurrences on islands. Combining information from 1,349 species lists and floras, A-Islands includes data on >6,500 plant species from 844 islands ranging in size from 18 m^2^ to 4,400 km^2^, exhibiting different degrees of isolation from the mainland, and spanning all major Australian climate zones. Of these, 251 islands have been repeatedly sampled up to 11 times making it possible to investigate temporal compositional change. A-Islands is open access and will be continuously updated. Its simple data structure consisting of three comma separated files allows easy integration with other Australian and global plant-occurrence databases and can serve as a repository for island research in Australia. Knowing which species occur on Australia′s islands will provide opportunities for future research, including studying changes in biodiversity and species-turnover within and among archipelagos, tests of classical island biogeography theory and as a baseline for ecological monitoring and conservation.

## Introduction

Continental islands serve as valuable models for studying the assembly of species into communities. Continental islands were once part of a continent but are now isolated by rising sea levels, typically located on shallow continental shelves (Whittaker et al. 2023). Within such island environments the processes and patterns of community assembly and change are comparatively easy to observe. Because the species-pools and communities in island ecosystems share similarities with those found on the mainland, but differ due to recurring colonization, speciation and extinction, thereby allowing the processes involved to be disentangled (MacArthur and Wilson 1967, Abbott 1977, Schrader et al. 2020, Schrader et al. 2023b). Islands also offer a model system to study the ecosystem consequences of fragmentation (Warren et al. 2015), a near-pervasive conservation concern in mainland habitats. Islands are also intrinsically valuable, harbouring high biodiversity that can include endemic species or important populations of threatened species (Kier et al. 2009, Fernandez-Palacios et al. 2021). Island endemic species also comprise a disproportionate extent of Australia’s and the world’s extinctions (Woinarski et al. 2019, Fernandez-Palacios et al. 2021). For all these reasons, islands are a top priority for conservation and present unrivalled opportunities for biodiversity research.

To formulate predictions and test hypotheses in island ecology, data sets that include a large number of islands across large geographic gradients are particularly useful. Large-scale approaches are important to pursue as community assembly processes are likely influenced by factors that vary along geographical gradients, such as prevailing dispersal vectors, soil, climate and degrees of island isolation and high replication is required to detect community dynamics (Sfenthourakis and Triantis 2009, Schrader et al. 2021, Schrader et al. 2023a). Further, investigating the dynamics of temporal turnover that offer insights into compositional change over time on islands – including testing influential theories such as the Equilibrium Theory of Island Biogeography (MacArthur and Wilson 1967, Schrader et al. 2023b) – requires datasets that hold information on repeatedly sampled islands.

Australia and its many surrounding islands provide a distinct opportunity to assemble a highly standardised database on the distribution of plants on islands. Australia’s coastline is fringed by >8,000 small-to medium-sized continental islands, which support considerable biodiversity (Moro et al. 2018). Many of Australia’s islands have been monitored extensively in the past seven decades thanks to the diligence of ecologists, botanists, and government agency staff (e.g., Abbott 1980, Harris et al. 2001, Batianoff et al. 2009).

Despite the depth of data on Australian island diversity (Kark et al. 2022), no comprehensive database combines plant species lists reported for islands with island properties such as area size and isolation. This hinders scientific advances in island biology. Also, many islands in Australia are severely affected by invasive species and land-use, but at the same time are potential safe havens for the conservation of threatened biota, crucial for protecting the many endangered plant and animal species of the continent from extinction (Legge et al. 2018, Bryant and Harris 2020, Woinarski et al. 2023). Practical conservation programs, such as island translocations, can benefit from information about local abiotic and biotic conditions on islands to improve success rates (Rayner et al. 2021). Predictions of suitability for potential invasive species can assist in guiding biosecurity prioritisation.

Here, we fill this gap for Australian islands and introduce a novel database – A-Islands – for plant occurrences on Australian islands. Islands considered for A-Islands were any landmass surrounded by ocean or coastal waters on the Australian shelf smaller than Tasmania. Oceanic islands, like Lord Howe or Norfolk Island, were not considered. Currently, A-Islands holds information on plant occurrence for 843 islands, which is over 10% of Australia’s c. 8,000 islands, spanning all major climate zones and biogeographical regions including 251 repeatedly sampled islands (Fig 1). Most of the islands included are small and of continental origin, where plant communities are maintained by immigration and extinction dynamics (Abbott and Black 1978, Schrader et al. 2023b). These islands allow for testing hypotheses on community assembly and spatio-temporal turnover dynamics.

**Fig. 1:**
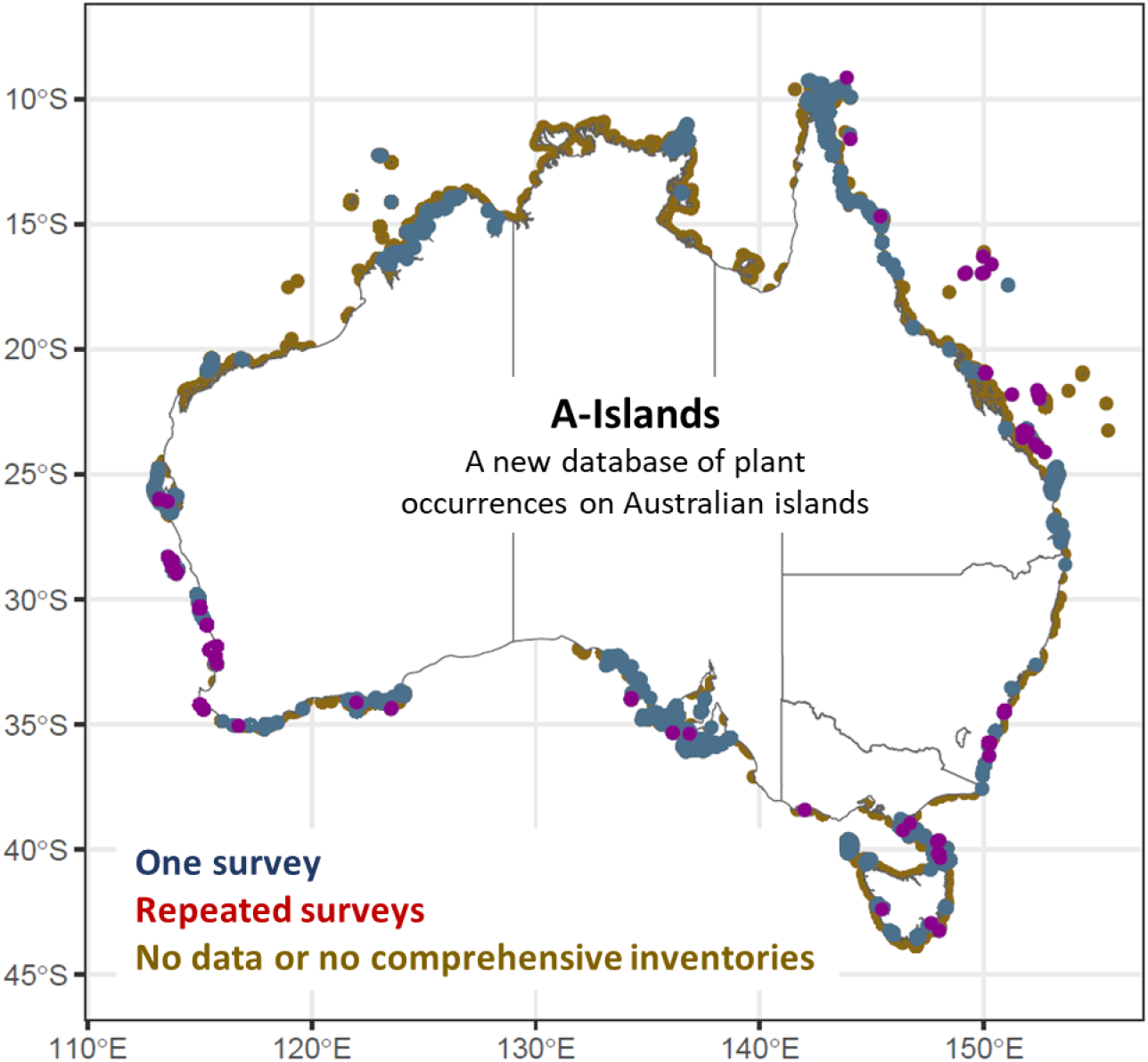
Island coverage in A-Islands, distinguishing between all islands that have been sampled once (blue; n=592) and sampled more than once (red; n=251). Brown are islands not included in A-Islands so far as they have not been (comprehensively) surveyed or no surveys were found. C. 10% of all Australian islands are included in the database spanning all major Australian climate zones.

We aim to continuously expand A-Islands with special focus on stimulating interest to resample island floras, as well as to conduct surveys on currently unsampled islands. Distinctive features of A-Islands are:

- Open access: all data in A-Islands are freely available, including species-by-island records and spatial island data. New versions will be released as new data accumulates.
- The focus on repeated surveys of island plant species: this is unique among island plant-occurrence databases.
- Standardised taxonomy: records are standardised according to national and global taxonomic references allowing for easy integration with other databases.

## Methods and Database Structure

To collate this database, we completed five major steps (Fig 2). These steps also serve as a framework for integrating new data and maintaining the database in the long run. The five steps consist of (1) searching for and access all relevant island survey information, (2) extracting species names and island information from the original references, (3) if available, providing information on species status (native vs naturalised) based on the original references, (4) updating and standardising species names and (5) allocating all islands to a consistent spatial shapefile layer including island coordinates and island area. All workflows are transparent and reproducible by linking to and retaining all original data (species names and status and island locations).

**Fig. 2:**
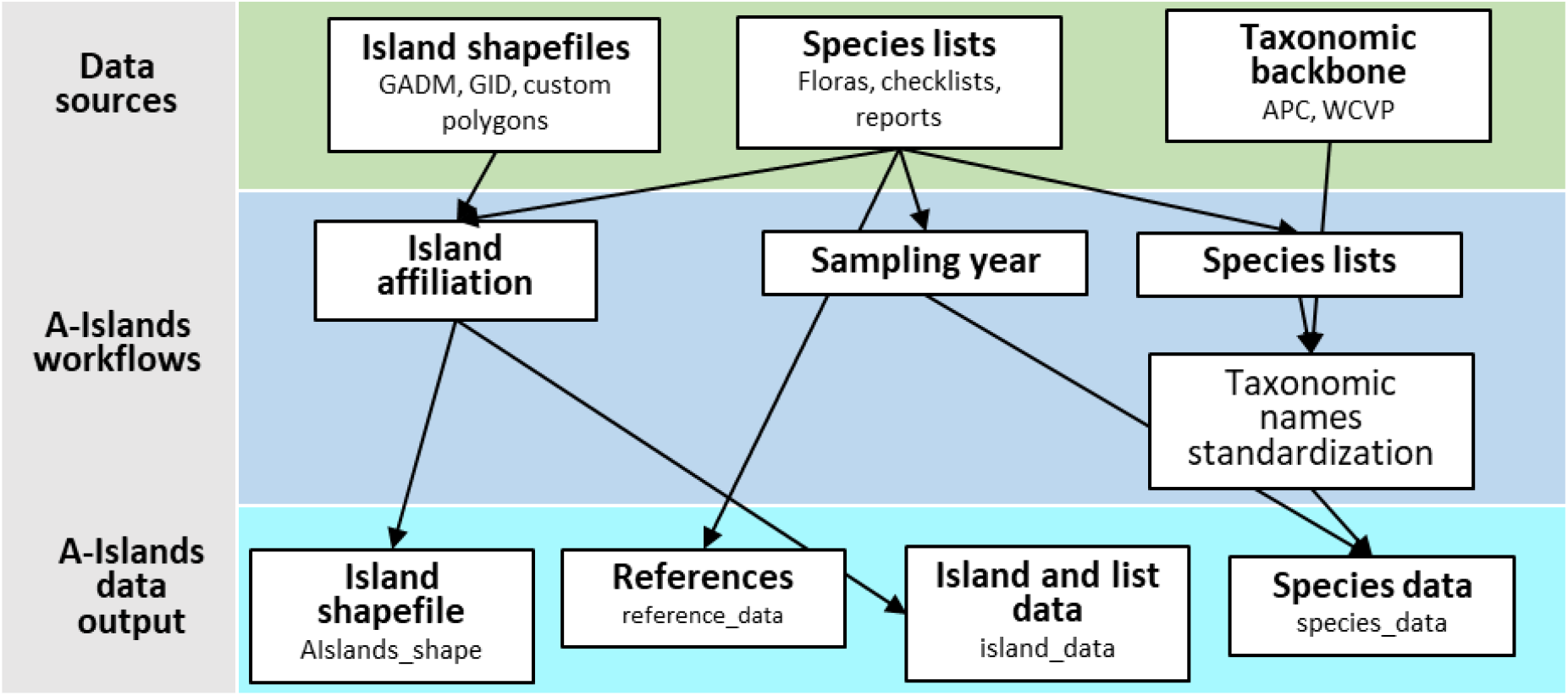
Workflows, data sources and data output in A-Islands.

1. **Searching for and access all relevant island survey information**: We conducted specific literature searches (Google Scholar, scanning of local Australian nature journals such as Western Australian Naturalist, Papers and Proceedings of the Royal Society of Tasmania or Atoll Research Bulletin) to detect and include as many island surveys as possible. Our search also included reaching out to researchers, botanists and environmental agencies who conducted island surveys in the past.
2. **Extraction of species names and island information:** We extracted species names and island-related information from the original references (Supplementary Table 1). For inclusion, only island species lists considered comprehensive by the original authors were used, thus excluding incomplete surveys or anecdotal species records. This approach enables A-Islands to facilitate biological analyses requiring thorough floristic accounts for individual islands. Many of these lists were compiled during specific field trips, with the database additionally providing the sampling year when available. Moreover, a number of islands have been sampled repeatedly, either by the same or by different authors, allowing for comparisons of species diversity over time. For each island, both the name provided in the original reference and, if applicable, a new name is included. In cases where some smaller islands lack official names, we used the IDs from the original references.
3. **Inclusion of species status information:** The majority of the original species lists included information about species status, indicating whether a species was native or naturalized. Some lists also contained details about species endemism. We retained and documented this information for each list, specifying whether species status was included, and for each species entry, whether it was categorized as native or naturalized. It is important to note that species status can be contentious or not conclusively resolved. Also, some species native to eastern Australia could, for example, be introduced on western Australian islands. We therefore recommend verifying the original inference as supplied by the authors of the species lists with authoritative and continuously updated lists, such as the Australian Plant Census (https://biodiversity.org.au; APC).
4. **Standardization of species taxonomy:** As species lists within A-Islands were compiled in various years, some dating back to the 1930s, differences in species taxonomy exist between records. To address this, we standardized the species taxonomy using two distinct taxonomic backbones. First, using the R package APCalign (Wenk et al. 2024), all species names were matched with the APC; this holds the most current and authoritative plant species taxonomy for Australia. The APC includes both currently accepted taxon names and outdated taxonomy (i.e. synonyms, basionyms, and misapplied names) and APCalign first aligns species names to any scientific name within the APC (correcting typos, standardising syntax), then updates the names to a currently accepted taxon concept. Updating names to those considered current by the APC facilitates seamless integration with other Australian plant databases, such as the trait database AusTraits (Falster et al. 2021) or the plot database HavPlot (Mokany et al. 2022), both of which also use the APC taxonomy. However, 615 taxa could not be matched with the APC. For these cases, we used the World Checklist of Vascular Plants (Govaerts et al. 2021). For 201 taxa that we could not match to either for the two taxonomies, we used the original name (mostly for species only identified to the genus or family level). We preserved all original species names from the references, allowing for ongoing taxonomic updates.
5. **Islands’ attributes, geospatial assignment, and archipelago allocation:** Each island was allocated a polygon within a GIS shapefile. This process involved cross-referencing with the Global Island Database (http://globalislands.net) and the Global Administrative Areas database (https://gadm.org). We checked visually whether Google Maps satellite image outlines corresponded with the polygons. Subsequently, we used the best-matching polygon from either the Global Island Database or the Global Administrative Areas. In cases where no suitable polygon existed or where there was poor correspondence with satellites’ shape and the available polygons, and any of the two shapefiles, we delineated a new polygon based on the satellite image. We then extracted island coordinates and island areas using the *sf* package in *R*. Island locations in the GIS shapefile allow for calculation of other geo-environmental variables such as isolation or perimeter or matching with climate and environmental raster layers. We also assigned all islands to archipelagos or island groups based on predefined archipelagos or shared attributes such as biogeographical region, substrate, or proximity. Recognizing potential ambiguity in archipelago delineation, we established two hierarchical archipelago levels. The first contains 13 archipelagos, encompassing large biogeographical regions like tropical Queensland or the entirety of the Bass Strait between mainland Australia and Tasmania. The second level comprises 35 archipelagos and provides finer-scale divisions, including smaller regions such as the Houtman Abrolhos archipelago in Western Australia or the Capricornia Cays in Queensland.

The output data are three data tables and one shapefile. The first data table (island_data) comprises island-level information, featuring a unique identifier for each island (island_ID), island names, island area (km^2^), longitude and latitude for the island centre, and the assigned archipelago (both levels 1 and 2). The second data table (species_data) encompasses all species (original and standardized names), their status (if applicable: native, endemic, naturalized), the corresponding island identifier (island_ID) for each island where a species was recorded and an ID for each list (list_ID; links to island_data and reference_data) a species was recorded in. The third data table (reference_data) provides information on the original references (linked via the list_ID to the species level data). These data tables are provided in comma-separated files (.csv). The shapefile includes polygons for each of the islands (linked by island_ID; note zthe geographic location of one island with species information could not be identified). Detailed information on these three data tables can be found in Tab. 1-3.

### Recorded Data

A-Islands contains species lists referring to 844 islands collated from 1,351 separate lists, with 251 islands (30%) having been sampled at least twice (Fig 3B). Of these 150 lists are linked to a specific sampling year and can be used for analyses on temporal change in species diversity. All lists in A-Islands originated from 134 references (Supplementary Table 1) and contain 59,773 species-by-island records. The number of unique original species names amounts to 7,711, which reduces to 6,291 after taxonomic standardisation at species-level, which is c. one fourth of all c. 24,000 Australian plant species (Chapman 2009). 45 islands only support a single species, whereas Kangaroo Island is the most species-rich island, with 1,160 species (Fig 3C). The islands are spread along the whole coastline of Australia with most islands (352) from Western Australia (Fig. 1). The smallest island included has an area of 18 m^2^ whereas K’gari (1,669 km^2^) and Kangaroo islands (4,400 km^2^) are the largest (Fig 3A; note that we treat Tasmania as part of the continent rather than as an island).

**Fig. 3:**
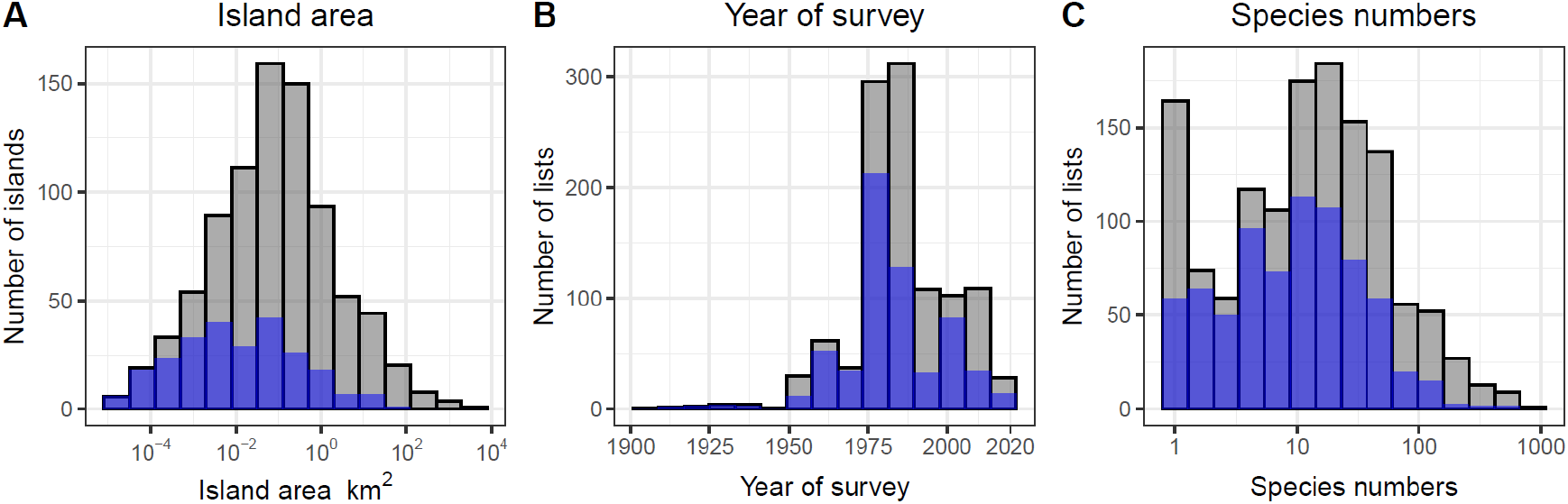
Quantity and distribution of data available in A-Islands for (A) island area (843 islands; geographic location needed to calculate area could not be identified), (B) year that a survey was conducted (1,108 island species lists) and (C) species richness (for 1,349 island species lists). Grey indicates total number of islands and lists and blue indicates islands that were repeatedly sampled and species lists from repeatedly sampled islands.

### Technical Validation

Technical validation consisted of three parts: (i) taxonomic, (ii) biogeographical, and (iii) geographical validation.

First, we validated the taxonomy by matching all species against the Australian Plant Census (https://biodiversity.org.au) and the World Checklist of Vascular Plants (Govaerts et al. 2021). Species names that could not be matched with at least one of these databases were checked manually for typographical and/or spelling errors and, if applicable, were corrected manually. This step was repeated until all species were regarded as correct (corrections were always checked against the original reference).

Second, we constructed a species-area relationship across all islands (Fig. 4). The z-values – a useful index for comparing species-area relationships (Matthews et al. 2016) – were 0.31 for all species, 0.28 for native species only and 0.12 for naturalised species only on the islands. The species-area relationship describes the increase in species richness with island area and is a strong pattern in ecology, holding true across nearly all island systems worldwide (Kreft et al. 2008, Matthews et al. 2019). We did this to visually test for outliers from the species-area relationship. Original references for these outliers were then checked to validate whether the species list was accurately transferred to A-Islands, and to assess whether the respective lists can be regarded as taxonomically comprehensive. We included only those lists to A-Islands that were considered comprehensive by the original author(s).

**Fig. 4:**
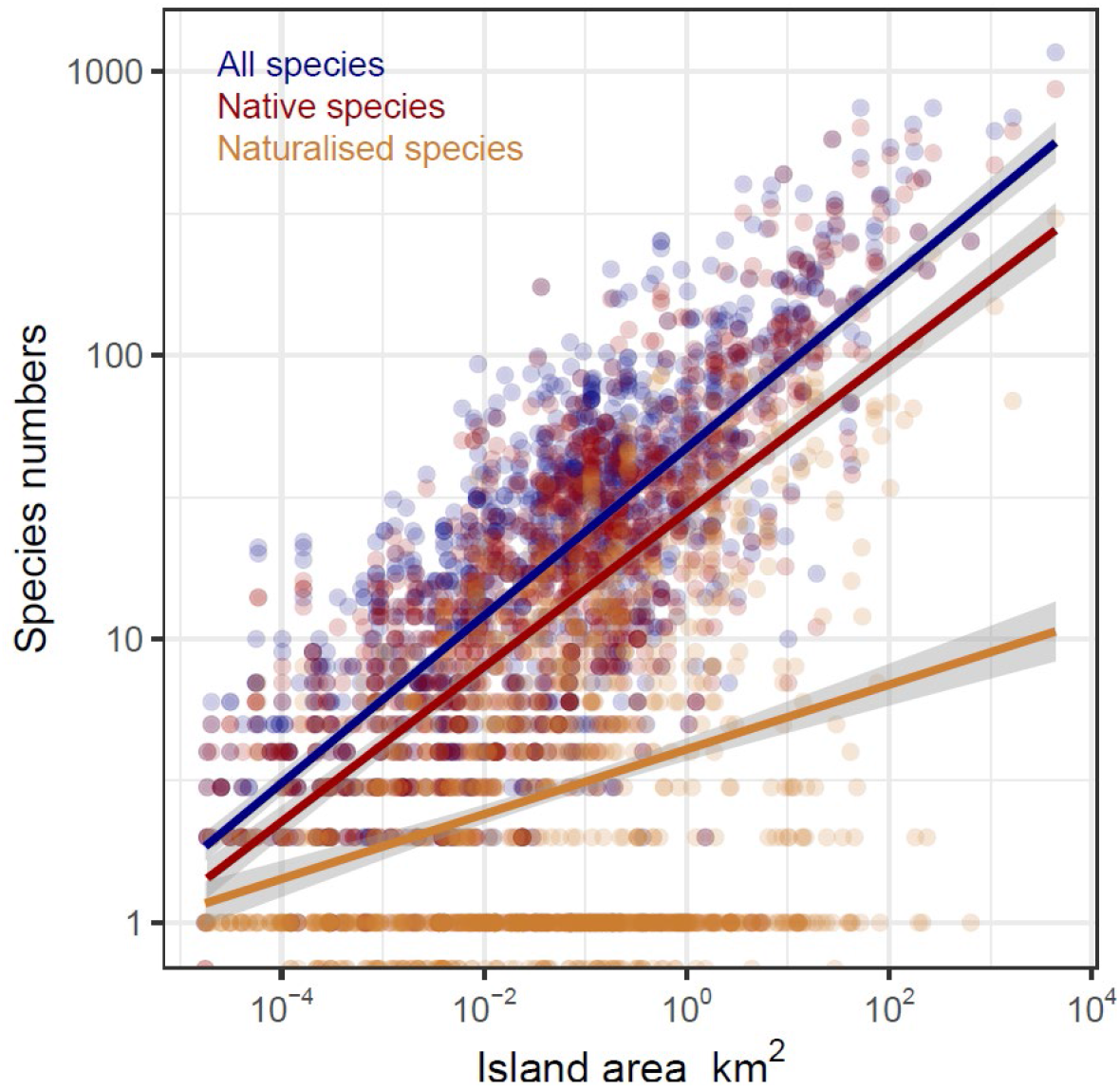
Species-area relationships for all 1,349 island surveys in A-Islands for native species (red), naturalised species (orange) and native and naturalised species combined (blue). Note that each point denotes a single sampling event on an island. Islands with >1 sampling event occur multiple times.

Third, we validated the geographical assignment of each island by testing whether islands covered in the same reference were spatially close together (all original references covered either a single island or two or more neighbouring islands).

### Usage Notes

A-Islands taxonomy aligns with the Australian Plant Census, which is Australia’s authoritative list of vascular plant taxonomy. This shared taxonomy facilitates integration with prominent Australian plant databases like AusTraits (Falster et al. 2021) or HavPlot (Mokany et al. 2022) which share this taxonomic backbone. Additionally, A-Islands can be effectively linked with the Global Inventory of Floras and Traits database (Weigelt et al. 2020), as both databases share a similar structure. Taxonomy can be updated using the function *create_taxonomic_update_lookup* from the *R* package *APCalign* (Wenk et al. 2024) (against the Australian Plant Census) and the *TNRS* package (Maitner and Boyle 2023) (against the World Checklist of Vascular Plants).

## Code Availability

All code used for maintaining and updating A-Islands will be made available upon request. Openly available R (v.4.2.1) packages and functions have been used for all major steps of data processing including taxonomic standardisation and spatial analyses (see methods for details).

## Acknowledgements

We are deeply indebted to all authors of the original references who visited or resided on the islands around Australia and made their surveys openly available in checklists, reports, and floras. This work was supported by a Macquarie University Research Fellowship grant to J.S. HK acknowledges funding of research unit FOR2716 DynaCom and Biodiversa+ BioMonI (533271599) from the German Research Foundation (DFG).

## Data Tables

**Table 1:** Information provided in data table reference_data. All references used are listed here including information whether native, naturalised and endemic (at the island level) species are indicated in the original reference.

| ref_ID | full_ref | native_indicated | naturalised_indicated | endemics_indicated |
| --- | --- | --- | --- | --- |
| Unique ID for each reference that links to island_data | Full reference | Includes information on native species (0, 1) | Includes information on naturalised species (0, 1) | Includes information on endemism at island level (0, 1) |

**Table 2:** Information provided in data table island_data. All islands and all sampling events are listed including information on island name, year of each species survey (if provided) coordinates, area and archipelago affiliation. Links to reference_data via ref_ID.

| ref_ID | list_ID | island_ID | island_name | survey_year | several_samplings | X | Y | area | arch_lvl_1 | arch_lvl_2 |
| --- | --- | --- | --- | --- | --- | --- | --- | --- | --- | --- |
| Links to reference_data | Unique ID for each species-island list | Unique ID for each island | Name of island | Year an island was surveyed (empty if not provided) | Indicates (0,1) if survey has been done in a single year. | X coordinates (longitude) of island | Y coordinates (latitude) of island | Area (km<sup>2</sup>) of island | Name of archipelago (level 1) | Name of archipelago (level 2) |

**Table 3:**
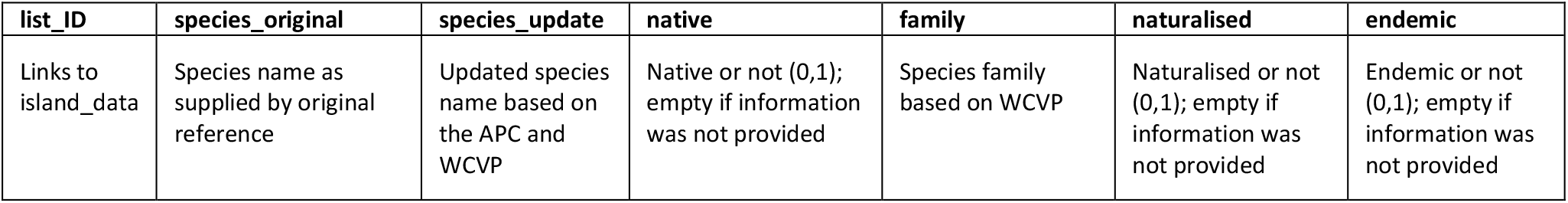
Information provided in data table species_data. All species from each list are shown here including their original name (as used in original reference), an updated name based on the most recent taxonomy and whether the species is native, naturalised or endemic on the respective island (if information provided). Links are to island_data via list_ID.

## Notes

**Conflict of interests** The authors declare no conflict of interests.

### Competing Interest Statement

The authors have declared no competing interest.

